# ASBAR: an Animal Skeleton-Based Action Recognition framework. Recognizing great ape behaviors in the wild using pose estimation

**DOI:** 10.1101/2023.09.24.559236

**Authors:** Michael Fuchs, Emilie Genty, Klaus Zuberbühler, Paul Cotofrei

**Author notes:** Contributing authors.

## Abstract

The study and classification of animal behaviors have traditionally relied on direct human observation or video analysis, processes that are labor-intensive, time-consuming, and prone to human bias. Advances in machine learning for computer vision, particularly in pose estimation and action recognition, offer transformative potential to enhance the understanding of animal behaviors. However, the integration of these technologies for behavior recognition remains underexplored, particularly in natural settings.

We introduce *ASBAR* (*Animal Skeleton-Based Action Recognition*), a novel framework that integrates pose estimation and behavior recognition into a cohesive pipeline. To demonstrate its utility, we tackled the challenging task of classifying natural behaviors of great apes in the wild.

Our approach leverages the OpenMonkeyChallenge dataset, one of the largest open-source primate pose datasets, to train a robust pose estimation model using DeepLabCut. Subsequently, we extracted skeletal motion data from the PanAf500 dataset, a collection of in-the-wild videos of gorillas and chimpanzees annotated with nine behavior categories. Using PoseConv3D from MMAction2, we trained a skeleton-based action recognition model, achieving a Top-1 accuracy of 75.3%. This performance is comparable to previous video-based methods while reducing input data size by approximately 20-fold, offering significant advantages in computational efficiency and storage.

To support further research, we provide an open-source, terminal-based GUI for training and evaluation, along with a dataset of 5,440 annotated keypoints for replication and extension to other species and behaviors.

All models, code, and data are publicly available at: https://github.com/MitchFuchs/asbar

## 1 Introduction

Direct observation and manual annotation of animal behaviors are labor-intensive, time-consuming, and prone to human error [1]. These methods also face significant limitations, such as information loss in low-visibility settings or during complex, fast-paced social interactions involving multiple individuals. Video recording and post-hoc annotation have thus become the preferred methods for studying animal behavior. They enable detailed identification and interpretation of behaviors, while also facilitating reliability testing and replication of coding. However, the manual annotation of videos remains a significant bottleneck, underscoring the need for automated systems that can streamline animal behavior analysis. Machine learning tools have the potential to identify relevant video sections containing social interactions and automatically classify behaviors, significantly expanding the scope and robustness of observational studies and enhancing our understanding of animal behaviors [2].

Recent advancements in machine learning and computer vision offer innovative avenues for building such systems. In particular, action recognition models can learn deep representations of video features and classify these features into behavior categories. Within deep learning, two primary approaches to action recognition have emerged: video-based methods and skeleton-based methods.

On one hand, video-based action recognition involves analyzing RGB video data to identify spatio-temporal patterns that characterize actions. This approach often relies on Convolutional Neural Networks (CNNs) [3] adapted to the temporal domain. Notable models include Two-Stream CNNs [4], C3D [5], I3D [6], (2+1)D ResNet [7], and SlowFast [8]. These methods have been extended to classify animal behaviors [9–14] and multimodal audio-visual data [15].

On the other hand, skeleton-based action recognition predicts behavior classes based on the skeletal structure and motion of the body [16, 17]. This approach relies on an additional preprocessing step, *pose estimation*, which detects body parts, such as joints and bones, and extracts their coordinates from video frames [18]. While skeleton-based methods require the added step of pose estimation, they offer several advantages for computational ethology [1, 2, 19]:

(i) *Cross-subject behavior recognition*: These models focus on skeletal motion rather than external appearance, allowing them to generalize across individuals within the same species (e.g., [20] for humans, [21, 22] for non-human animals).
(ii) *Robustness to visual setting changes*: Video-based models are sensitive to lighting conditions, background variations, and other subtle changes in input data [23–25]. Comparatively, skeleton-based methods are less affected by these variations [26].
(iii) *Reduced computational complexity*: Extracting pose coordinates reduces the dimensionality of video data, lowering computational costs and power consumption [16]. This is particularly beneficial for field researchers with limited resources.
(iv) *Geometric quantification*: Pose estimation provides a pre-computed geometric representation of body motion and behavior [1, 27].

A major challenge for skeleton-based methods in animal behavior analysis lies in obtaining accurate pose-estimated data. While human pose estimation benefits from extensive open-source datasets and state-of-the-art detectors, achieving similar performance for animals remains challenging. Fortunately, there has been a surge in annotated animal pose datasets (e.g., Animal Kingdom [28], Animal Pose [29], AP-10K [30], OpenMonkeyChallenge [31], OpenApePose [32], MacaquePose [33], Horse-30 [34]) and open-source pose estimation frameworks (e.g., DeepLabCut [35–37], SLEAP [38, 39], AniPose [40]). Despite these advancements, the use of pose estimation for behavior recognition remains underexplored, particularly in natural settings. Challenges include the lack of datasets with both keypoint coordinates and behavior annotations, and the tendency to treat pose estimation and behavior recognition as separate tasks rather than parts of an integrated approach.

To address these challenges, we introduce ASBAR, an innovative framework for animal skeleton-based action recognition. Our contributions include:

- An integrated pipeline, which combines a DeepLabCut-based pose estimation module with a behavior recognition module from MMAction2 [41]. The pipeline is encapsulated in a terminal-based GUI, allowing researchers to train and evaluate models without programming knowledge, even in remote or cloud-based environments.
- A robust primate keypoint detector, by leveraging the OpenMonkeyChallenge dataset [31], which spans 26 primate species. Additionally, we provide detailed performance metrics for species and individual body parts and release 5,440 high-quality keypoint annotations for great apes in their natural habitats.
- A methodology for wild behavior analysis using the PanAf500 dataset [11], a two-hour collection of camera trap videos annotated with nine locomotive behaviors. We demonstrate that our skeleton-based pipeline achieves performance comparable to existing video-based methods.

## 2 The ASBAR Framework

ASBAR is an integrated data and model pipeline (marked in red in Fig 1) designed to address two sequential machine learning tasks: *pose estimation* and *action recognition*.

**Fig. 1:**
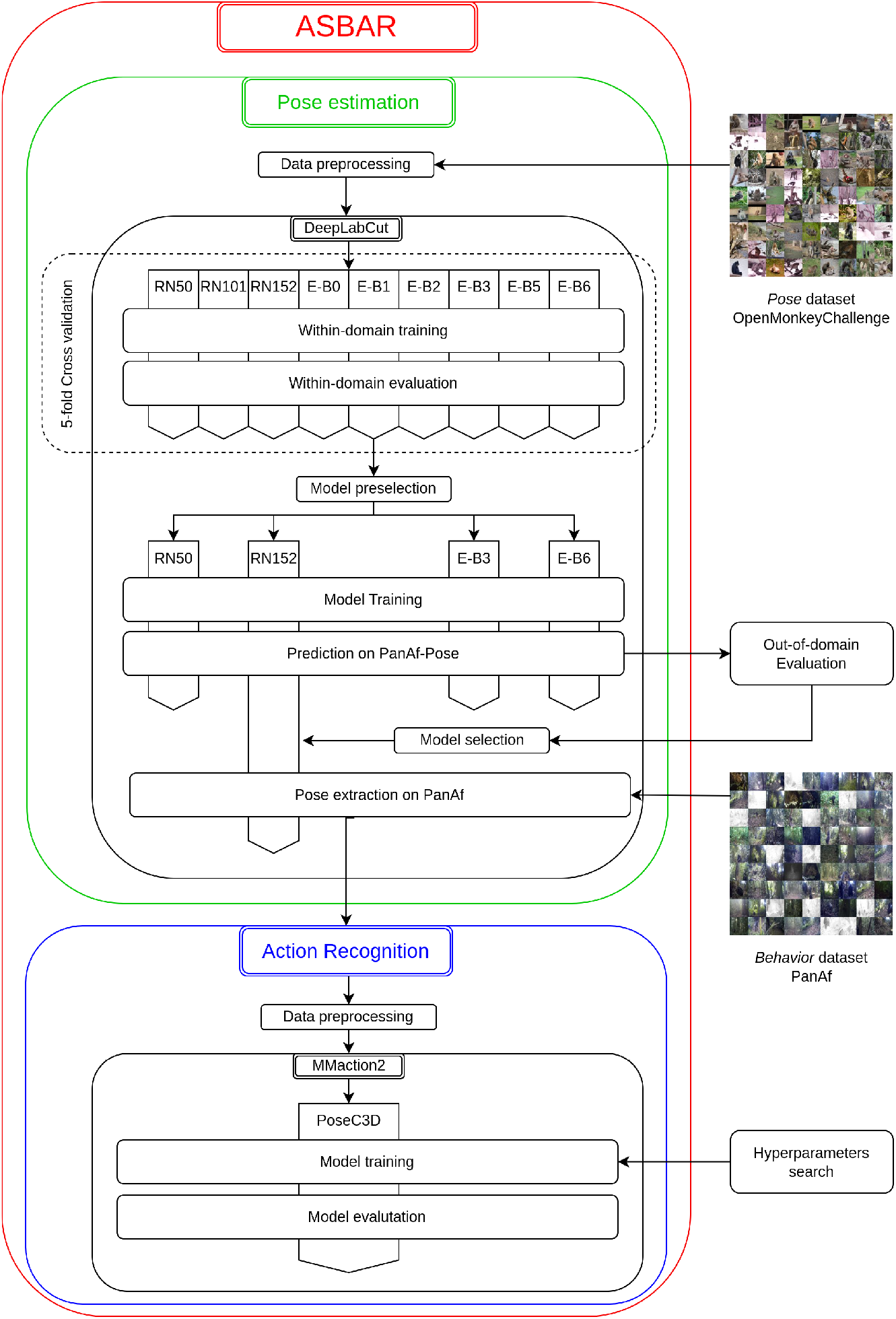
The ASBAR Framework. The ASBAR framework’s data and model pipeline (red) comprises two modules: a *pose estimation* module (green) based on DeepLabCut and an *action recognition* module (blue) integrating models from MMAction2.

The first module, responsible for animal pose estimation (marked in green in Fig 1), incorporates key features of DeepLabCut (DLC), a widely used framework for multianimal, markerless pose estimation [35–37]. This module includes functionality for project creation, dataset preparation, model training and evaluation, configuration editing, and video analysis.

The second module, focused on behavior recognition (marked in blue in Fig 1), integrates APIs from MMAction2 [41], a comprehensive platform for action recognition and video understanding. Specifically, this module employs PoseConv3D [17], a convolutional neural network (CNN) model tailored for skeleton-based action recognition.

To demonstrate the capabilities of the ASBAR framework for animal skeleton-based behavior recognition, we selected a particularly challenging task for our experiments: classifying natural great ape behaviors in the wild.

### 2.1 Pose and Behavior Datasets

ASBAR operates on two distinct datasets: a *pose* dataset and a *behavior* dataset. The pose dataset contains images annotated with 2D keypoint coordinates and is used to train the pose estimator model. The behavior dataset comprises video clips annotated with specific behaviors.

Ideally, both datasets originate from the same visual distribution—for example, pose images being a subset of video frames from the behavior dataset. However, in practice, annotating a dataset with both pose and behavior labels is time-consuming and costly. A pragmatic approach involves combining pose and behavior datasets from different visual distributions. For instance, Internet images annotated with keypoints can complement video data labeled with behaviors recorded in the wild. In such cases, the pose dataset is considered *within-domain*, while the behavior dataset is referred to as *out-of-domain*.

### 2.2 Pose Estimation Module

The task of estimating the coordinates of anatomical keypoints, denoted as *pose estimation*, is a crucial prerequisite for skeleton-based action recognition. In most cases, each RGB frame of a video clip of an action is preprocessed to extract the set of (*x, y*) coordinates in the image plane of each keypoint and its relative confidence *c*. In practice, this transformation is often performed via supervised CNN-based machine learning models trained for keypoint detection (Fig 2).

**Fig. 2:**
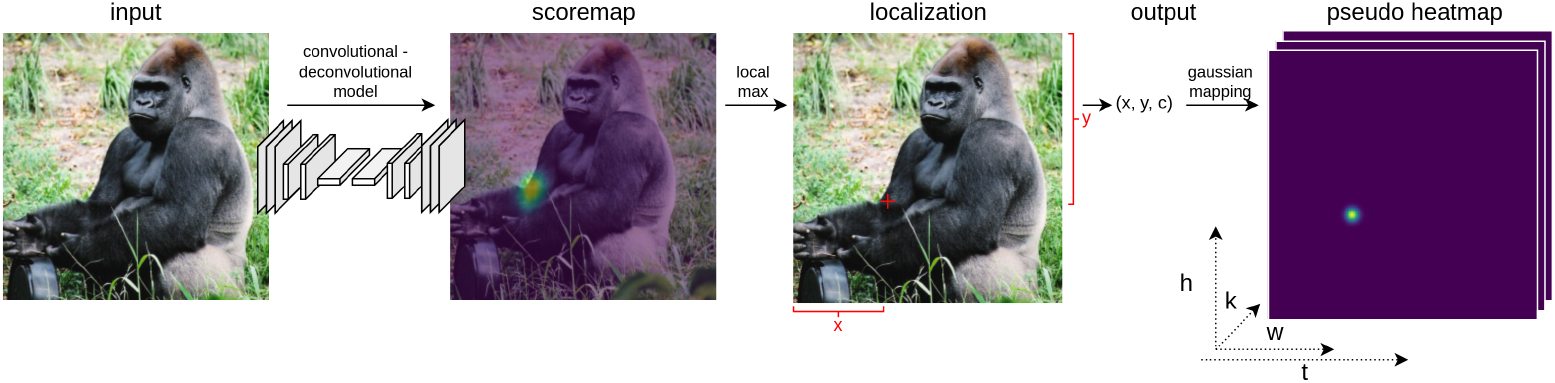
From RGB image to pseudo-heatmaps. The transformation of an RGB image into a 3D heatmap volume. An input image is passed through a Conv-Deconv architecture to output a probabilistic scoremap of the keypoint location (e.g., the right elbow). By finding a local maximum in the scoremap, the location coordinates and confidence can be extracted. Using a Gaussian transformation, a pseudo heatmap is generated for each keypoint and used as input of the subsequent behavior recognition model.

The goal of the pose estimation module is to extract pose information from the behavior dataset using a trained pose estimator. This module encompasses four key functionalities: *data preprocessing, model benchmarking, model selection*, and *pose extraction*.

#### Data Preprocessing

The module provides four preprocessing steps:

- *Data formatting*: Ensures the pose dataset meets DeepLabCut’s structural requirements.
- *Data selection*: Allows users to customize the dataset by selecting specific species, annotated keypoints, or excluding invisible keypoints. For example, users can limit the dataset to chimpanzees and bonobos, focusing on three visible keypoints (e.g., eyes and nose).
- *Dataset splitting*: Supports options for no cross-validation, 5-fold cross-validation, or 10-fold cross-validation to enable statistical validation of the model’s performance.
- *Configuration setup*: Enables customization of DeepLabCut’s configuration files, including training hyperparameters.

#### Model Benchmarking

Given the importance of high pose prediction performance for behavior recognition accuracy, the framework facilitates benchmarking various pose estimation models. Users can evaluate models with different backbones (e.g., ResNet or EfficientNet) and depths (e.g., ResNet50, ResNet101, EfficientNet-B0).

#### Model Selection

When the pose and behavior datasets share the same visual distribution, benchmarking results suffice for model selection. However, for out-of-domain behavior datasets (Sect.2.1), additional evaluation is necessary, as within-domain performance does not guarantee robustness to visual domain shifts. Models with EfficientNet backbones, for example, have demonstrated superior generalization to out-of-distribution scenarios compared to ResNet models [34]. Evaluating out-of-domain performance involves comparing model predictions with manually labeled video frames from the behavior dataset. More details are provided in Sect.3.3.

#### Pose Extraction

The selected model extracts pose information from the behavior dataset. Users can specify a particular model snapshot or allow the framework to choose the snapshot with the lowest test set error.

### 2.3 Action Recognition Module

Skeleton-based action recognition involves the classification of a specific action (performed by human or non-human individuals) from a sequence of skeletal joint data (i.e., coordinate lists), captured by sensors or extracted by markerless pose estimators (Fig. 3).

**Fig. 3:**
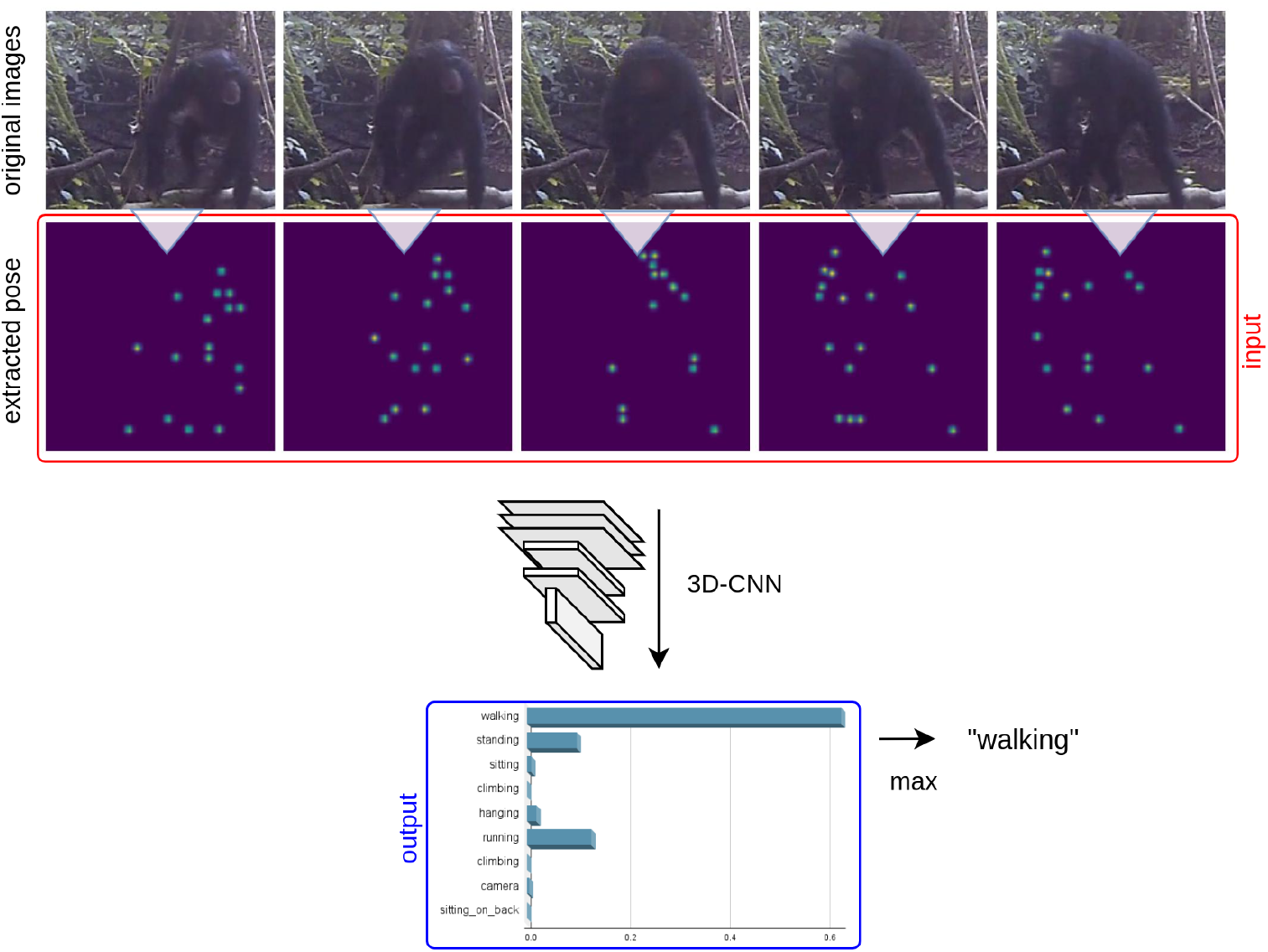
From extracted poses to behavior classification. From a set of consecutive RGB frames (e.g., 20 in our experiments), the animal pose is extracted, transformed into pseudo-heatmaps, and stacked as input of the behavior recognition model. A 3D-CNN is trained to classify the represented action into the correct behavior category (e.g., here ‘walking’)

The action recognition module classifies behaviors in the behavior dataset using pose data extracted from the first module. This module includes three functionalities: *data preprocessing, model training*, and *model evaluation*.

#### Data Preprocessing

To enable behavior recognition, the module implements four preprocessing steps:

- *Prediction filtering*: Retains only the highest-confidence keypoint predictions for each frame. For each keypoint, the most confident coordinates within the labeled bounding box are kept, while others are discarded.
- *Data sampling*: Extracts sequences of consecutive frames that meet specific time and behavior label constraints (see Sect. 3.5 for details).
- *Data formatting*: Converts skeleton data into a structure compatible with PoseConv3D.
- *Configuration setup*: Allows customization of PoseConv3D’s configuration, including hyperparameter settings.

#### Model Training

Users can train several variations of PoseConv3D available in the MMAction2 toolbox [17, 41]. These variations include different 3D-CNN backbones (e.g., SlowOnly [42], C3D [5], X3D [8]) with an I3D classification head [6]. Training can be distributed across multiple GPUs to reduce computation time.

#### Model Evaluation

The module produces probabilistic classifications, returning a ranked list of behavior candidates with associated confidence probabilities. The behavior with the highest confidence is used to calculate Top-1 Accuracy, the percentage of correctly predicted samples. Other metrics, such as Top-*k* Accuracy (percentage of ground-truth behaviors within the top-*k* predictions) and Mean Class Accuracy (average Top-1 accuracy across behavior classes), are also supported.

## 3 Materials and Methods

### 3.1 Datasets and Data Annotation

For the classification of great ape behaviors in their natural habitat, we utilized two primary datasets: OpenMonkeyChallenge and PanAf500. Additionally, we manually labeled a subset of keypoint coordinates from PanAf500, referred to as PanAf500-Pose.

#### OpenMonkeyChallenge

OpenMonkeyChallenge (OMC) [31] is a benchmark dataset containing 111,529 images of 26 primate species, designed for non-human primate pose estimation challenges. The dataset includes images sourced from the web, three U.S. National Primate Research Centers, and multiview cameras at the Minnesota Zoo. Each image is annotated with species, bounding box coordinates, and 2D pose information for 17 keypoints, including the nose, eyes, head, neck, shoulders, elbows, wrists, hips, tail, knees, and ankles. For occluded keypoints, annotators were instructed to provide the most likely location and specify visibility.

The dataset is divided into training (60%), validation (20%), and testing (20%) subsets. While the testing annotations are withheld for competition purposes, we combined the training and validation sets to create a comprehensive *pose* dataset containing 89,223 images. Examples of these images are shown in Fig 4 (left).

**Fig. 4:**
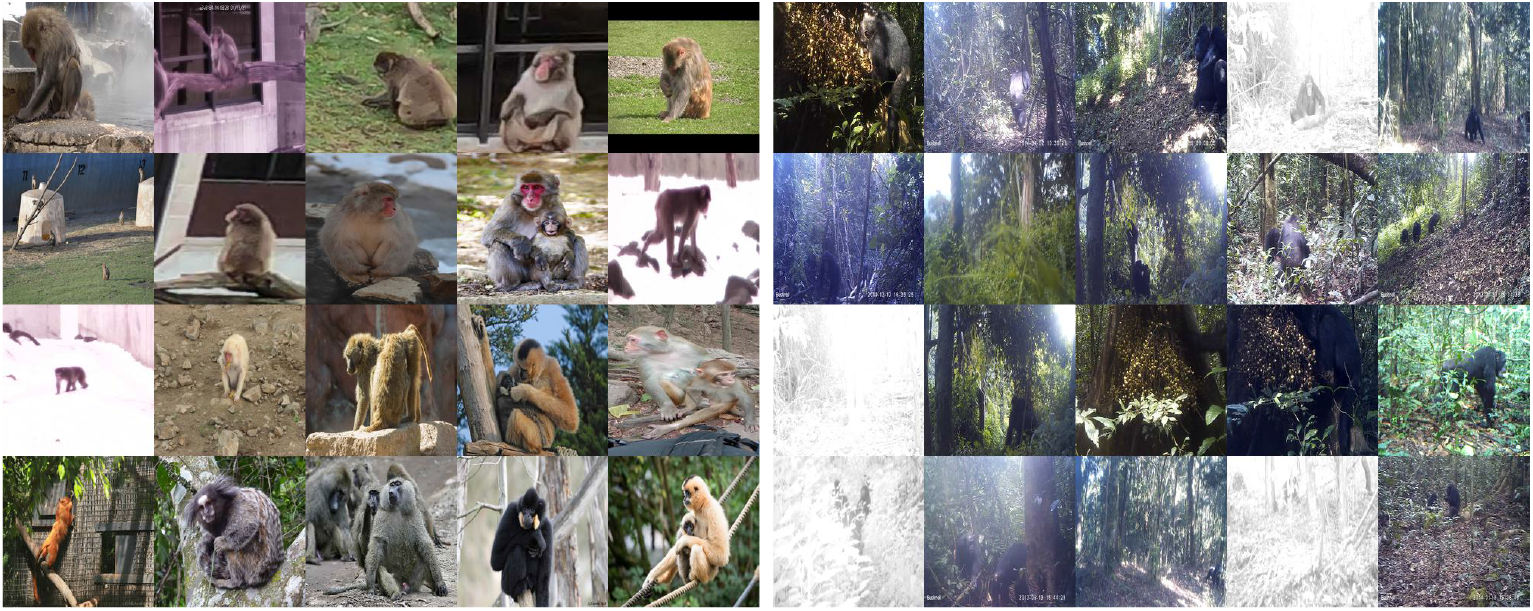
Examples from the *pose* and *behavior* datasets. (*Left*) Sample images from the *OpenMonkeyChallenge* dataset, one of the largest collections of primate images annotated with 2D poses. This dataset contains over 100,000 images from 26 primate species. (*Right*) Sample video frames from the *PanAf500* dataset, comprising 500 videos of gorillas and chimpanzees recorded in African forests using camera traps. The dataset includes annotations for bounding boxes and behaviors. Visual challenges include small individual sizes due to camera distance, abundant vegetation, nocturnal imaging, and varying backgrounds.

#### PanAf500

The Pan African Programme “The Cultured Chimpanzee” [43] aims to enhance understanding of the ecological and evolutionary factors influencing chimpanzee behavioral diversity. This program has amassed thousands of hours of footage from camera traps deployed in Central African forests. The PanAf500 dataset consists of 500 15-second videos (180,000 frames at 24 FPS) of chimpanzees and gorillas, annotated with bounding box coordinates for ape detection [44, 45] and behaviors for action recognition [11]

The dataset includes nine annotated behaviors: ‘walking,’ ‘standing,’ ‘sitting,’ ‘running,’ ‘hanging,’ ‘climbing up,’ ‘climbing down,’ ‘sitting on back,’ and ‘camera interaction.’ The class distribution exhibits a long-tail pattern [46], with three *head* classes (‘walking,’ ‘standing,’ and ‘sitting’) each containing over 1,000 samples. In contrast, *tail* classes such as ‘running,’ ‘climbing up,’ ‘climbing down,’ ‘sitting on back,’ and ‘camera interaction’ have fewer than 100 samples each. Examples from this dataset are displayed in Fig 4 (right).

#### PanAf500-Pose

To supplement the PanAf500 dataset, we manually annotated 5,440 keypoints across 320 images, using the same keypoints as in OMC. The annotation process involved three steps:

- *Image selection*: We first shortlisted 4,000 images using predictions from ResNet152 and EfficientNet-B6 models based on high overall prediction confidence (Section 3.3). From this shortlist, 320 frames were manually selected to represent diverse scenes, lighting conditions, postures, sizes, and species, while minimizing consecutive frames;
- *Mini-clip generation*: For each selected frame, we generated a 34-frame mini-clip (24 frames before and 10 frames after) to capture motion and aid in labeling occluded keypoints;
- *Keypoint annotation*: We employed a semi-automated annotation process, initially leveraging predictions from the ResNet152 model. These predictions were refined using DeepLabCut’s Napari plugin [47]. In the first phase, a non-trained annotator (MF) adjusted the predictions. The annotations were then finalized by a great ape behavior and signaling expert (EG) [48], ensuring high-quality labels.

### 3.2 Evaluation Metrics

#### 3.2.1 Evaluation Metrics for Pose Estimation

##### Mean Average Euclidean Error (MAE)

MAE is the primary evaluation metric in DeepLabCut and measures the average Euclidean distance between the ground-truth labels 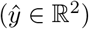 and the model predictions (*y* ∈ ℝ^2^):

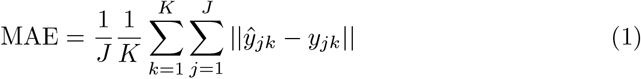

Here, *J* is the number of images (e.g., 89,223 in OMC) and *K* is the number of keypoints (e.g., 17 in OMC). Refer to Fig. A1 in Appendix A for a visual comparison of predictions and MAE examples.

##### Percentage of Correct Keypoint - nasal dorsum (PCK)

PCK measures the percentage of keypoints that fall within a specified distance of the ground-truth. PCK is computed as:

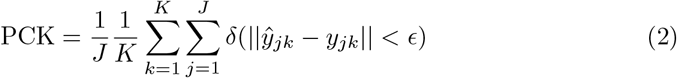

Here, *δ*(·) is an indicator function that outputs 1 when the condition is met and 0 otherwise. The threshold distance *ϵ* is equal to the nasal dorsum length, defined as the distance between the midpoint of the eyes and the tip of the nose calculated for each frame as:

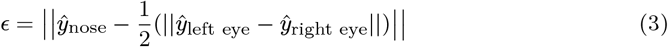

See Fig. B2 in Appendix B for an example of nasal dorsum calculation.

##### Normalized Error Rate (NMER)

NMER quantifies the mean normalized error by dividing the raw pixel distance between the predicted and ground-truth keypoints by the square root of the bounding box area [34]:

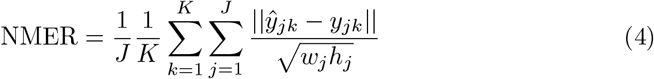

Here, *w* and *h* denote the width and height of the bounding box, respectively.

Both PCK and NMER are size- and distance-normalized metrics, unlike MAE, making them more robust for evaluating diverse scenarios.

#### 3.2.2 Evaluation Metrics for Action Recognition

Similar to [11], we use the three following action recognition metrics:

##### Top-1 Accuracy

Top-1 Accuracy measures the percentage of samples where the model’s highest-confidence prediction matches the ground-truth label.

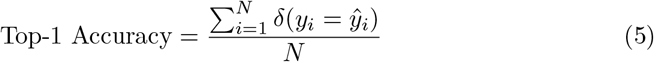

Where,

- *N* : Total number of predictions/instances
- *y*_*i*_: True class for the *i*-th instance
- 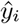 : Predicted class for the *i*-th instance (the class with the highest confidence score)
- *δ*(·): Indicator function, returns 1 if the condition is true, 0 otherwise

##### Top-3 Accuracy

Top-3 Accuracy measures the percentage of samples where the ground-truth label appears within the top three predictions of the model.

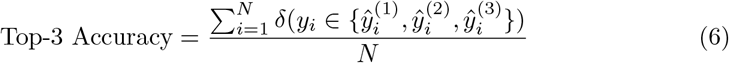

Where,

- 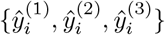 : set of top-3 predicted classes for the *i*-th instance, ranked by confidence scores

##### Mean Class Accuracy (MCA)

MCA calculates the average accuracy across all classes, giving equal weight to each class irrespective of sample size. See Fig. 10 for details.

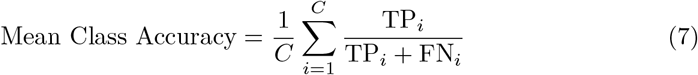

Where,

- *C*: Total number of classes
- TP_*i*_: True positives for class *i* (number of instances from class *i* correctly predicted as class *i*)
- FN_*i*_: False negatives for class *i* (number of instances from class *i* not predicted as such)

### 3.3 Methods for Pose Estimation

#### Data Preprocessin

For our experiment, we utilized all annotated data from the OpenMonkeyChallenge (OMC) dataset, encompassing 26 species and 17 keypoints, including invisible ones. Due to OMC’s large size, a 5-fold cross-validation approach was chosen for model benchmarking.

#### Within-domain Models Benchmarking

We evaluated the within-domain performance of nine pose estimation models, including three ResNet architectures (RN-50, RN-101, RN-152) and six EfficientNet variants (B0, B1, B2, B3, B5, B6) [49, 50]. All models were pretrained on ImageNet [51] and then trained on OMC for 40,000 iterations, a duration estimated as sufficient for loss convergence in preliminary tests using the largest network (EfficientNet-B6).

Default training hyperparameters and augmentation settings were used except for the batch size, which was set to 16 (the maximum fitting into the GPU memory for EfficientNet-B6 on an NVIDIA A100 40GB). The learning rate schedule followed the defaults: 1.0e-04 until iteration 7,500, 5.0e-05 until iteration 12,000, and 1.0e-05 for the remainder. We performed 5-fold cross-validation, splitting the dataset into 80% training and 20% testing subsets, ensuring that all 89,223 images were included in the test set once. All models were trained remotely on the Google Cloud platform with NVIDIA A100 (40GB) or V100 (16GB) GPUs, using ASBAR’s GUI (see Fig. D4 in Appendix D for examples of elements).

Model snapshots were saved every 5,000 iterations and evaluated on the test set. Each model’s eight snapshots were evaluated across all five folds (8× 5 = 40 evaluations per model). To handle the computational load, we customized DLC’s evaluation pipeline to batch data processing and limit evaluations to the test set. While this modification was not integrated into the released framework, the default DLC evaluation method remains available in ASBAR’s GUI for reproducibility.

#### Model Shortlisting

Since the ultimate goal was to predict keypoints on the out-of-domain *behavior* dataset, we shortlisted four models based on their robustness and generalization potential: (i) ResNet-152: Best-performing within-domain model; (ii) ResNet-50: Widely used and popular among researchers; (iii) EfficientNet-B6: Demonstrated strong generalization to out-of-domain data in prior studies [34]; (iv) EfficientNet-B3: Despite lower within-domain accuracy, it balances strong out-of-domain generalization [34] with low computational cost (1.8G FLOPs versus 4.1G for ResNet-50 and 11G for ResNet-152).

The shortlisted models were retrained on the full OMC dataset (89,223 images) with no test set. Training was extended to 100,000 iterations to accommodate the larger training dataset. All other hyperparameters were unchanged. Model snapshots were saved every 5,000 iterations, producing 20 snapshots per model for evaluation.

### 3.4 Methods for Pose Extraction

After evaluating pose estimation performance (see Sect. 4.1 for results), ResNet-152 was selected for pose extraction. This model was applied to all frames in the *behavior* dataset to predict keypoint candidates.

Given the visual differences between the *pose* and *behavior* datasets, the pose estimation model’s confidence threshold was lowered to 10^*−*6^ to maximize keypoint candidate generation and minimize cases of “no prediction.” Skeletal poses were extracted by filtering these candidates to retain only the 17 keypoints with the highest confidence scores within the annotated bounding boxes.

### 3.5 Methods for Behavior Recognition

#### Data Preprocessing

The methodology proposed by [11] was followed for data sampling. A minimum threshold of 72 consecutive frames (equivalent to 3 seconds) exhibiting the same behavior was set to ensure the inclusion of prolonged and meaningful behavioral patterns. Selected video clips were divided into samples of 20 consecutive frames, with no gaps or overlaps between samples.

The dataset was randomly split at the video level into training, validation, and testing subsets, using a 70-15-15 distribution. The pose extraction output was formatted and stored as triplets of (*x, y*, confidence) coordinates for each keypoint.

#### Model Training

A PoseConv3D model [17] with a ResNet3dSlowOnly backbone and an I3D classification head was selected for behavior recognition. This architecture was chosen for its strong performance on NTU60-XSub [20], a benchmark dataset for human action recognition, as reported by [17].

To adhere strictly to a skeleton-based approach, the model was trained exclusively on pose-estimated data, without incorporating the multimodal RGB+Pose capability. Only joint keypoints (excluding limbs) were used, with a sigma value of 0.6. Keypoint confidence scores were not considered, given the low confidence threshold during pose extraction.

The model weights were initialized from pretraining on the FineGym dataset [52]. Training was conducted for 50 epochs using two NVIDIA RTX 2080 Ti GPUs (2 × 11GB). A class-balanced focal loss [46] was employed to address the imbalanced class distribution (*β* = 0.992, *γ* = 2). Other hyperparameter choices included: batch size of 32; initial learning rate of 0.005 (with momentum of 0.9 and cosine annealing); weight decay of 0.01 and a dropout ratio of 0.8 (i.e. strong regularization to avoid overfitting). Other hyperparameters and augmentation settings followed those used in [17].

## 4 Results

This section details the experimental results. First, we present the evaluation of pose estimation models, including both within-domain and out-of-domain results, leading to the selection of an optimal model for pose extraction (Section 4.1). Additional insights into model performance at keypoint and species levels are provided in Section 4.2. Finally, skeleton-based behavior classification results are reported and compared to existing studies (Section 4.3).

### 4.1 Results of Pose Estimation

#### Within-domain evaluation

We compared the performance of all nine models after 40,000 iterations by constructing 95% confidence intervals for MAE using a t-distribution (*α* = 0.025, *ν*= 4), given the small sample size from cross-validation. The results (as seen in Figure 5) show that: (i) ResNet-152 achieved the best performance (14.05 ± 0.199), statistically outperforming other models; (ii) ResNet-101 ranked second (14.334 ± 0.080); (iii) EfficientNet-B5 (14.958±0.299), EfficientNet-B6 (14.981 ± 0.288), and ResNet-50 (15.098 ± 0.12) had overlapping confidence intervals, making their performances statistically indistinguishable; (iv) EfficientNet-B3 (15.455 ± 0.097) and EfficientNet-B2 (15.519 ± 0.25) performed slightly worse; (v) EfficientNet-B1 (16.031 ± 0.167) and EfficientNet-B0 (16.631 ± 0.546) exhibited the highest error rates.

**Fig. 5:**
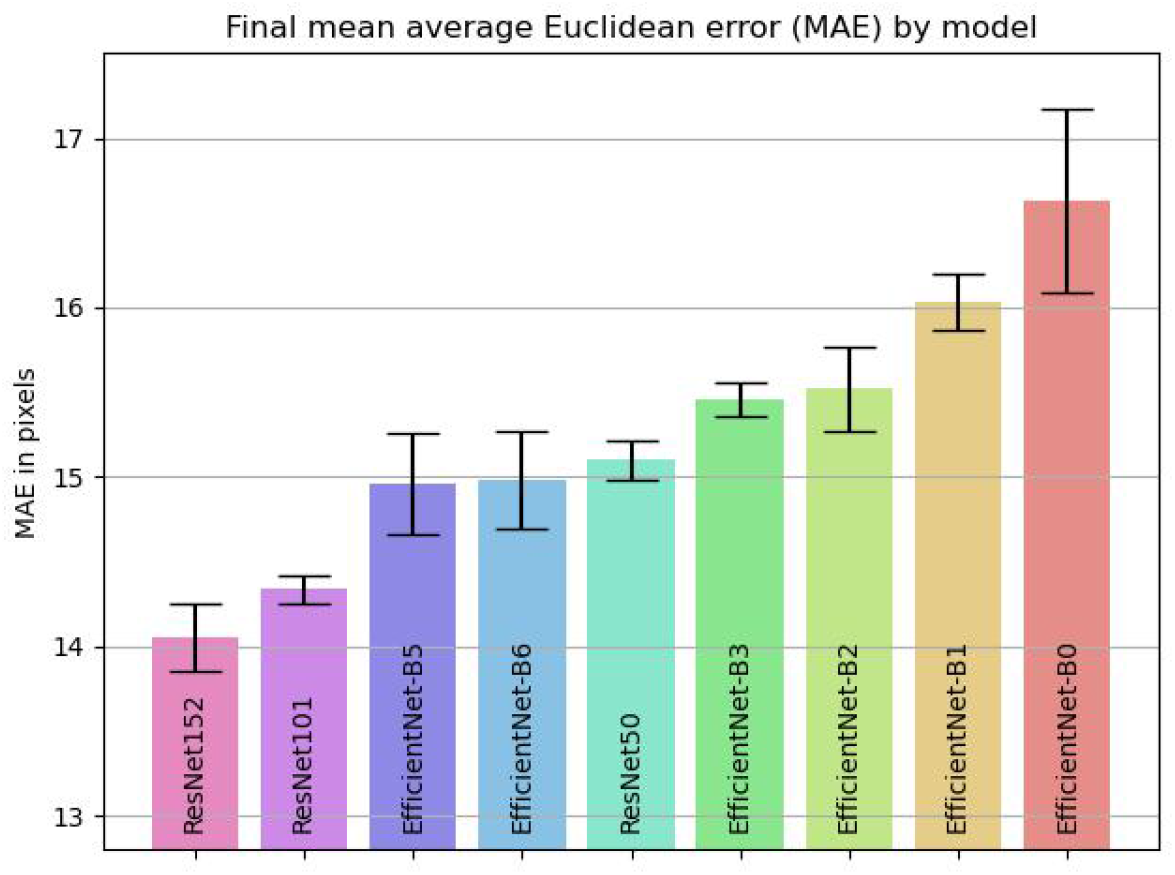
Final within-domain model performance. Mean and 95% confidence intervals of the MAE (in pixels) after 40,000 iterations (end of training). Disjoint confidence intervals indicate statistically significant differences. ResNet-152 demonstrates significantly better performance compared to all other models in this task.

In addition, the performance of each model during training is visualized through deviation charts of their snapshot variants, showing the mean and standard deviation of MAE and PCK in Figure 6.

**Fig. 6:**
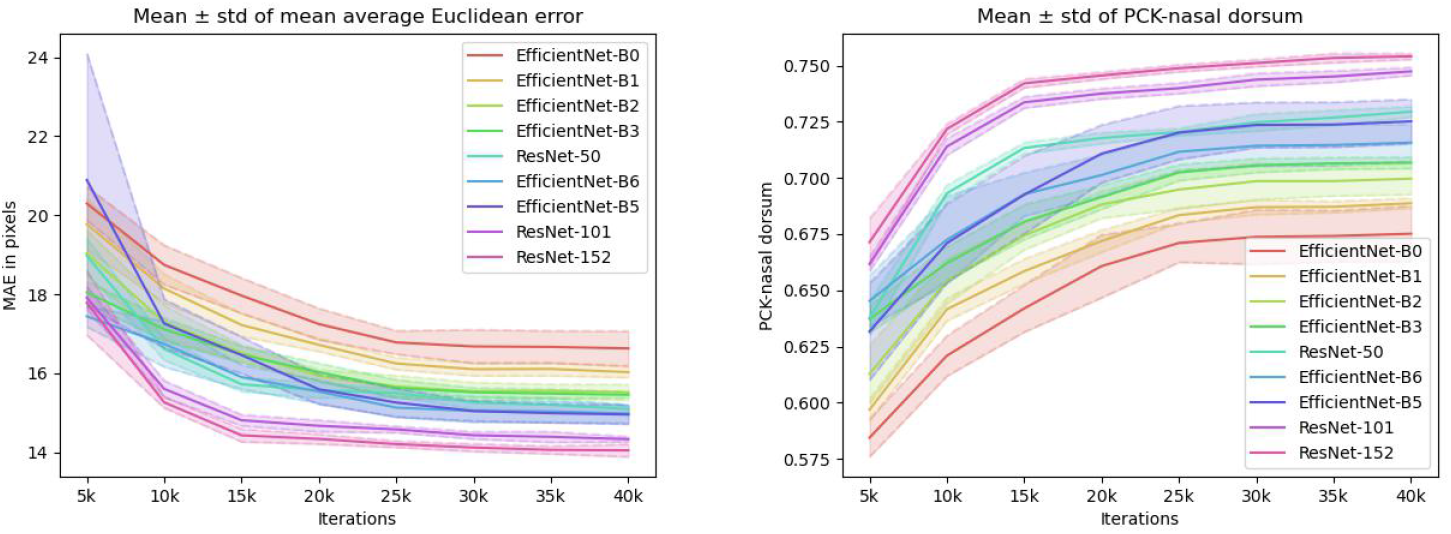
Model’s relative performance throughout ‘within-domain’ training. The mean ± std of the Mean Average Euclidean Error (MAE) in pixels (*left*, lower is better) and percentage of correct keypoint (PCK nasal dorsum) (*right*, higher is better) for all nine model variations. Evaluation results of 5-fold cross-validation on test set data, at every 5,000 iterations.

#### Out-of-domain Evaluation

Each saved model snapshot was tested on the PanAf500-Pose ground-truth annotations. To reduce the influence of noisy predictions, the minimum confidence threshold for pose prediction was maintained at the default value of 0.1.

Our results indicate that ResNet-152 generalizes best to out-of-domain data (Fig 7), achieving the highest overall PCK-nasal dorsum of 54.17% across all keypoints (*n* = 5, 440) and the lowest normalized error rate (NMER) of 10.19%.

**Fig. 7:**
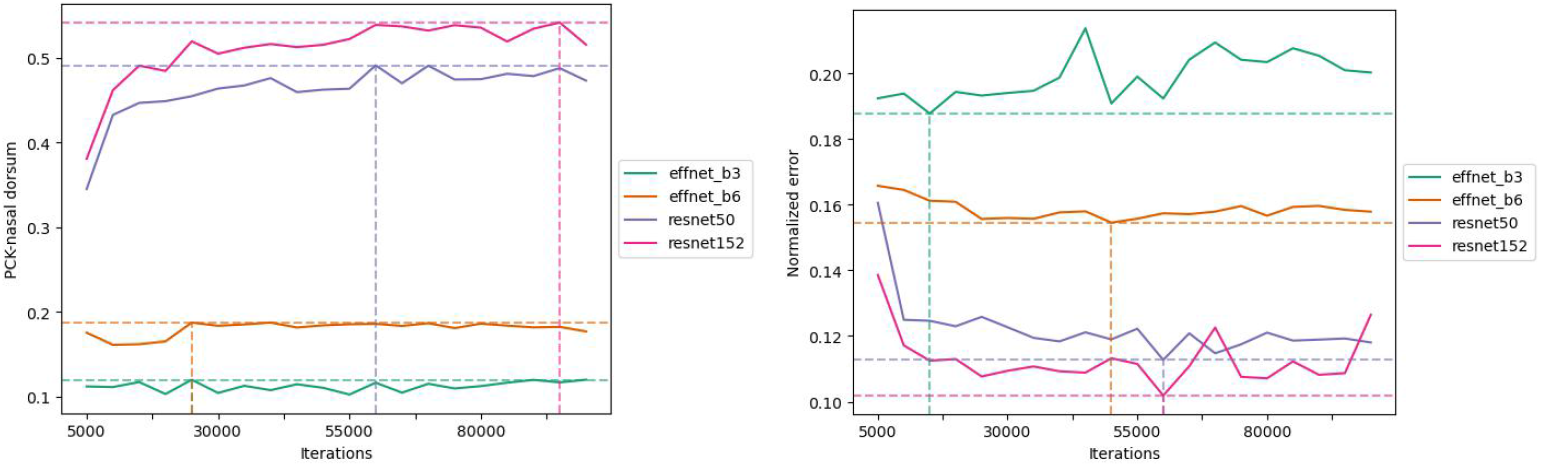
Out-of-Domain performance on PanAf500-Pose. Models are evaluated using two metrics that account for the animal’s relative size and distance: PCK nasal dorsum (*left*, higher is better) and normalized error rate (*right*, lower is better). ResNet-152 demonstrates superior performance in predicting great ape poses in their natural habitat. Vertical and horizontal dashed lines indicate the maximum and minimum values, along with the corresponding number of iterations. ResNet-152 at 60,000 iterations is selected for pose extraction.

#### Pose extraction

Based on ResNet-152’s performance at 60,000 iterations—achieving a detection rate of 53.9% (very close to the highest observed value) and a minimal NMER of 10.19%—this snapshot was selected as the final keypoint detector for pose extraction.

### 4.2 Alternative Performance Evaluation

To gain deeper insights into the final pose estimation model’s performance, we evaluated it across keypoints and species using both OMC and PanAf500-Pose datasets. A confidence threshold of 0.1 was applied throughout.

#### 4.2.1 Keypoint Detection Rate

Our analysis revealed that not all keypoints are equally detectable for non-human primates. Detection rates, computed as the cumulative distribution of predicted distances in pixels (Fig 8), highlight the following trends at test time for OMC (*n* = 89, 223 × 17 = 1, 516, 791):

**Fig. 8:**
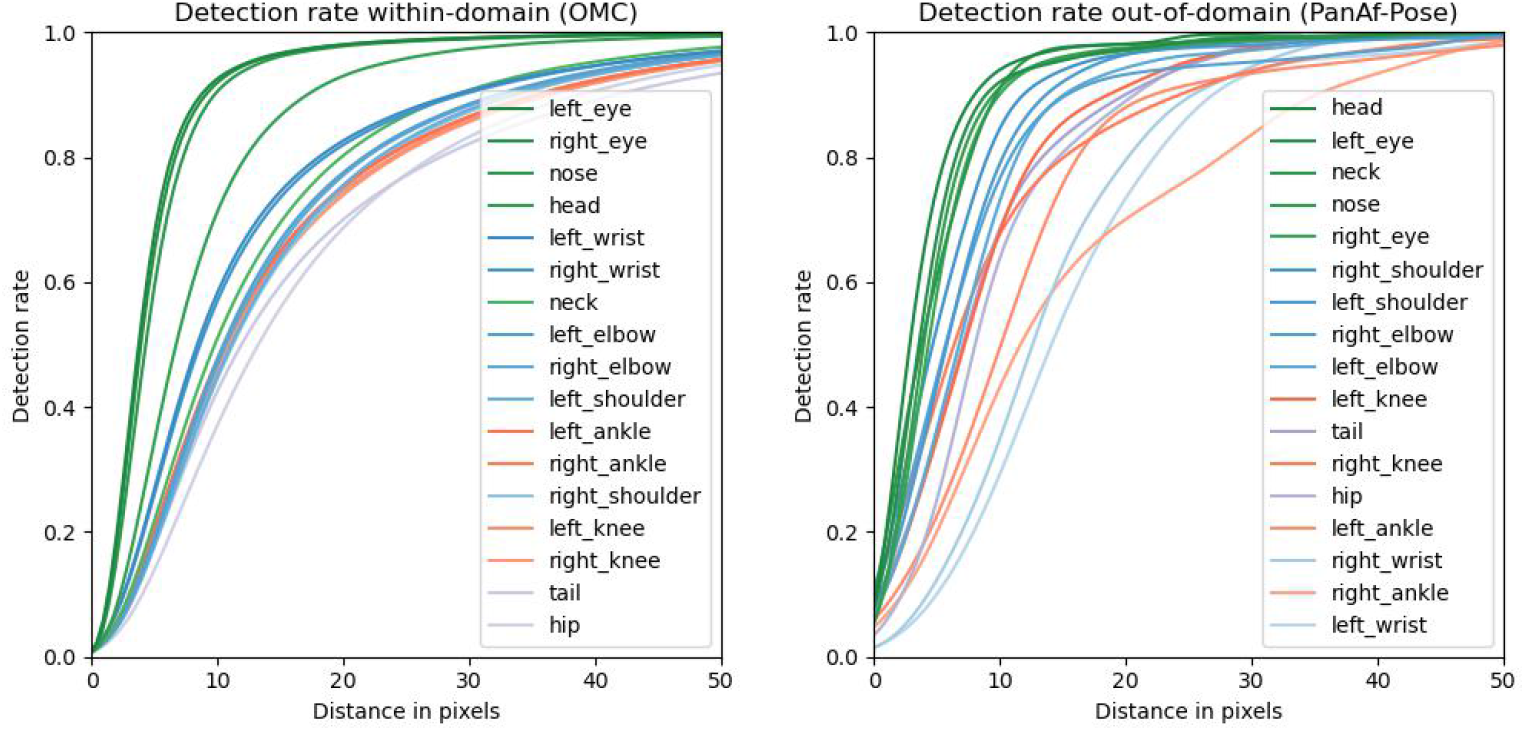
Keypoint detection rate on within-domain vs. out-of-domain test data. The keypoint detection rate, defined as the percentage of keypoints detected within a given pixel distance, is shown for OMC (*left*) and PanAf500-Pose (*right*). For example, within a distance of 10 pixels or less, the nose is detected in approximately 95% of the 89,223 images in OMC. In contrast, the tail is detected within the same distance in only about 38% of cases.

- Facial features (e.g., nose, eyes) are the easiest to detect.
- Keypoints on the head are more accurately predicted than those below the neck.
- Upper body limbs (e.g., wrists, elbows, shoulders) are detected more reliably than lower body limbs (e.g., ankles, knees).
- Limb extremities (e.g., wrists, ankles) are predicted more accurately than proximal keypoints (e.g., elbows, knees).
- Hip and tail positions are the most challenging to predict accurately.

These trends can be attributed to (i) the distinct visual features of facial keypoints, (ii) the prominence of heads and limb extremities, and (iii) the occlusion and ambiguity of lower body parts and tails.

Comparing results from PanAf500-Pose (*n* = 320 × 17 = 5, 440) shows a similar S-shaped distribution, indicating the model’s robustness to domain shifts. However, lower detection rates for specific keypoints may result from the precise annotations in PanAf500-Pose compared to OMC, where annotations are occasionally inconsistent (e.g., labeling fingers instead of wrists or toes instead of ankles).

#### 4.2.2 Per-Species Accuracy

To evaluate species-specific performance, we analyzed chimpanzees (*n* = 6, 190) and gorillas (*n* = 1, 777) in OMC using the normalized error rate (NMER) and 95% confidence intervals (Fig 9).

**Fig. 9:**
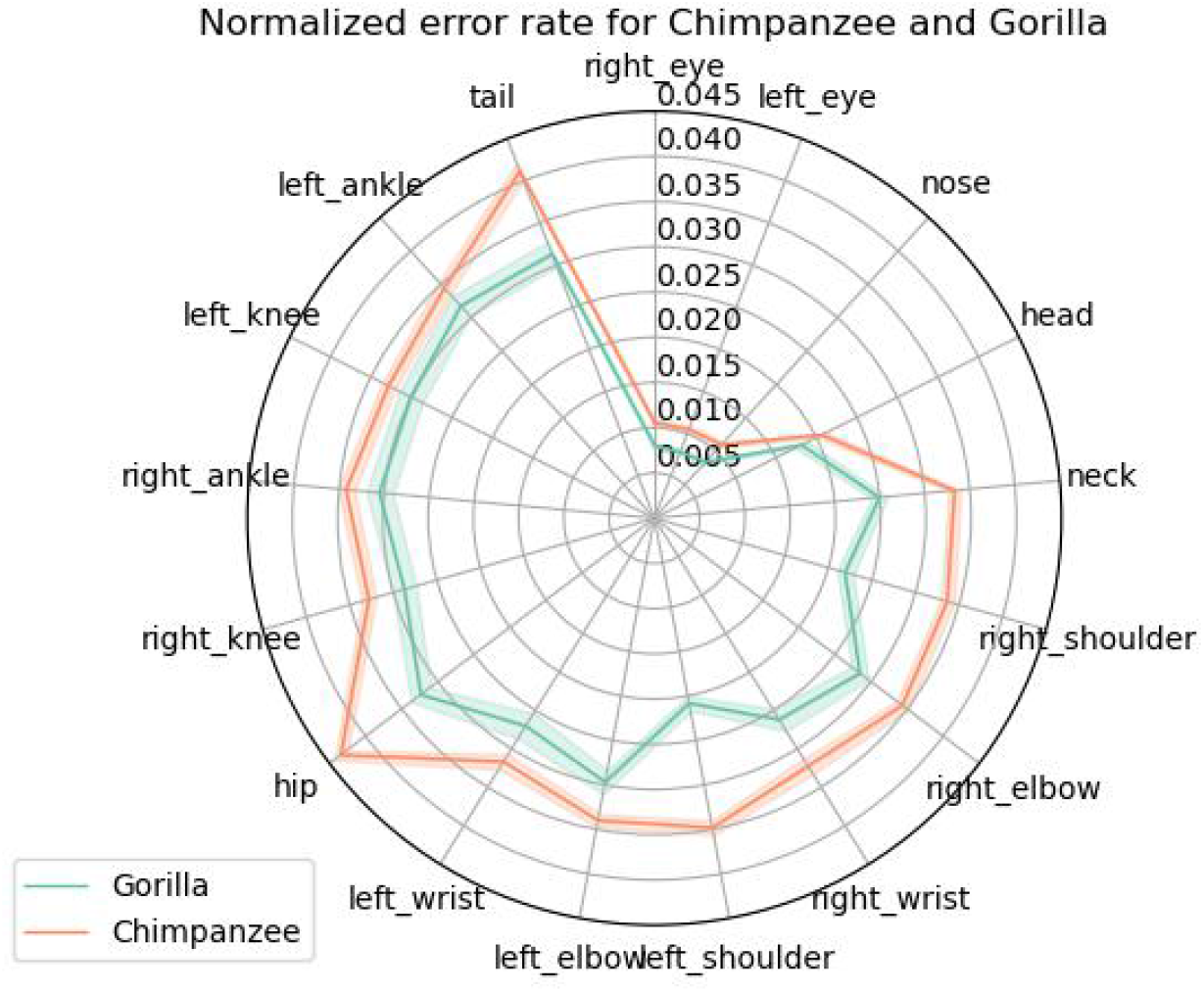
Normalized error rate for chimpanzees and gorillas in OMC. Mean and 95% confidence intervals for the normalized error rate (NMER). Disjoint confidence intervals indicate statistical significance. The model demonstrates lower error rates for all gorilla keypoints, suggesting higher prediction accuracy for this species.

**Fig. 10:**
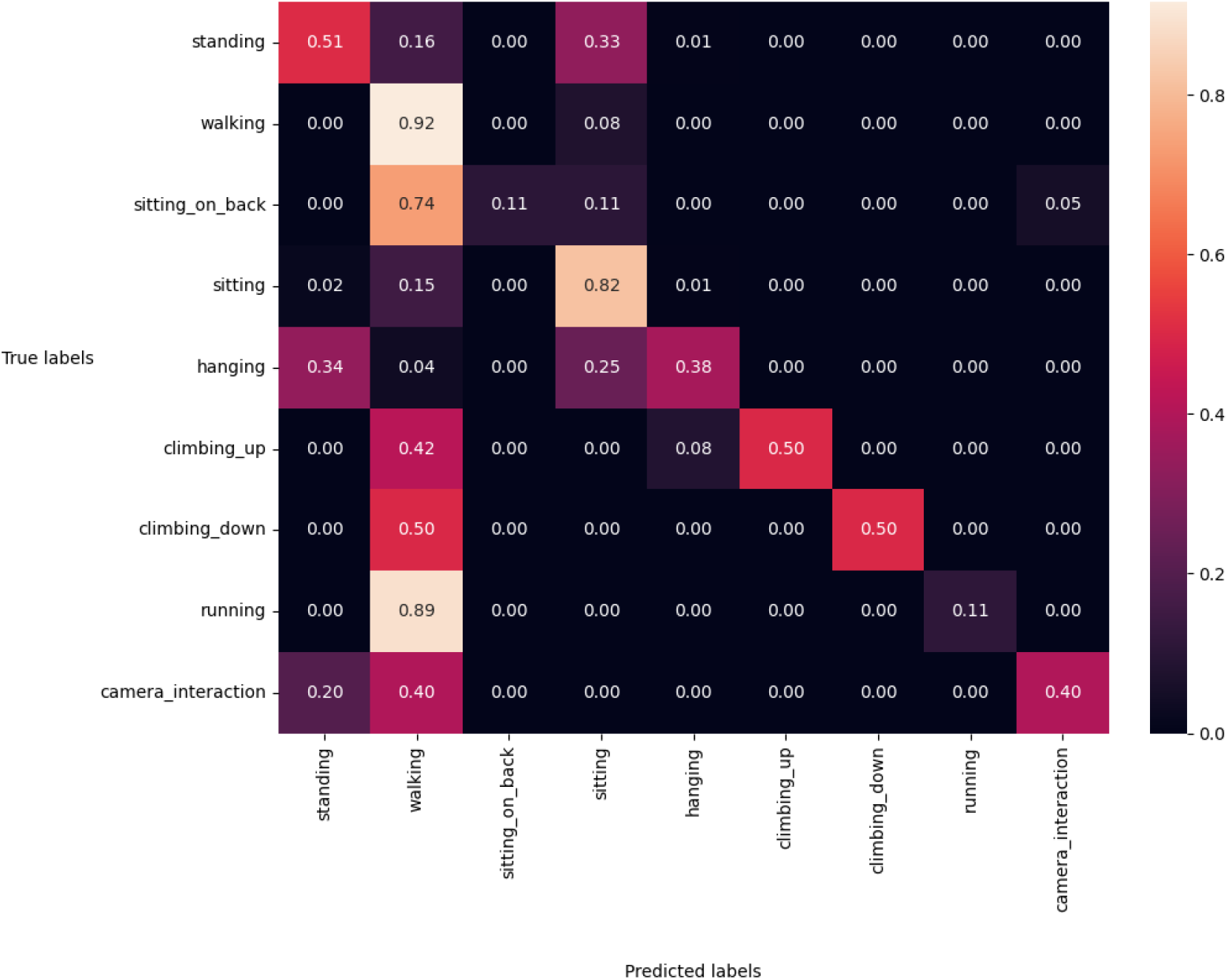
Normalized confusion matrix of behavior recognition. For each true behavior label (rows), the percentage of predictions across all predicted behaviors (columns) is shown. For instance, 51% of samples labeled as ‘standing’ were correctly classified, while 16% were misclassified as ‘walking’ and 33% as ‘sitting.’ The diagonal cells represent the per-class accuracy, and their average corresponds to the Mean Class Accuracy (MCA) metric. A perfect classification model would yield a normalized confusion matrix with values of 1 on the diagonal and 0 elsewhere.

Results indicate a statistically significant dependence on species, with gorillas consistently showing lower error rates than chimpanzees across all keypoints. This suggests that the model detects keypoints more accurately for gorillas. Additional species-level analysis is provided in Fig C3 in Appendix C.

### 4.3 Results of Behavior Recognition

The results of behavior classification, summarized in Table 1, demonstrate the successful application of our skeleton-based action recognition pipeline for animals. In the context of automating the recognition of great ape behaviors in the wild—a highly relevant yet challenging task—our approach achieves accuracy comparable to other video-based techniques, such as those reported in [11].

**Table 1:** Performance comparison with previous studies. Comparison of Top-1 Accuracy, Top-3 Accuracy, and Mean Class Accuracy (MCA) between ASBAR and previous video-based methods. ASBAR achieves comparable performance to video-based approaches across all metrics.

|  | Approach | Top1 Accuracy | Top3 Accuracy | MCA |
| --- | --- | --- | --- | --- |
| Two-Stream CNNs [11] | video-based | 73.5% | 94.1% | 42.3% |
| ASBAR | skeleton-based | **75.3%** | **95.4%** | **47.0%** |

To the best of our knowledge, this is the first use of a skeleton-based method for classifying great ape behaviors. Notably, the entire behavior dataset after pose extraction (i.e., the input features for the behavior classifier) requires less than 60 MB of storage in text format—approximately 20 times smaller than the storage requirements of the same dataset using a video-based approach. For ethologists working in the field, where computational, storage, and transfer resources are often limited, this represents a significant improvement without sacrificing performance in behavior recognition.

The normalized confusion matrix of the final behavior recognition model is shown in Fig.10. The model tends to overfit on *head* behavior classes, which have more samples in the dataset (see Sect.2.1). For instance, the model frequently overpredicts ‘walking,’ the second most represented class, at the expense of *tail* classes. The false positive rates (i.e., misclassification rates) for ‘walking’ on ‘sitting on back,’ ‘climbing up,’ ‘climbing down,’ ‘running,’ and ‘camera interaction’ are 0.74, 0.42, 0.50, 0.89, and 0.40, respectively.

Interestingly, the other two *head* classes—’standing’ and ‘sitting’—show near-zero false positive rates for the same *tail* classes. This discrepancy may be explained by the static nature of ‘standing’ and ‘sitting,’ which involve stationary poses, compared to the dynamic movements in ‘walking’ and most *tail* classes, where pose estimation accuracy may be lower.

Additionally, the true positive rates for ‘sitting on back’ and ‘running’ (i.e., their per-class accuracy) are extremely low, at 0.11 each. Both are predominantly misclassified as ‘walking.’ This likely stems from the similarity in skeletal poses across these behaviors, making it challenging for the model to differentiate between them using only skeleton data, particularly given the limited sample sizes for these classes.

## 5 Discussion

Despite the growing availability of open-source resources, such as large-scale animal pose datasets and machine learning toolboxes for pose estimation and human skeleton-based action recognition, their integration for animal behavior recognition—particularly in natural settings—remains largely unexplored. With ASBAR, a framework combining animal pose estimation and skeleton-based action recognition, we provide a comprehensive data and model pipeline, methodology, and GUI to assist researchers in automatically classifying animal behaviors via pose estimation. We hope these resources will become valuable tools for advancing the understanding of animal behavior within the research community.

To illustrate ASBAR’s capabilities, we applied it to the challenging task of classifying great ape behaviors in their natural habitat. Our skeleton-based approach achieved accuracy comparable to previous video-based studies for Top-K and Mean Class Accuracies. Additionally, by reducing the input size of the action recognition model by a factor of approximately 20 compared to video-based methods, our approach requires significantly less computational power, storage space, and data transfer resources. These qualities make ASBAR particularly suitable for field researchers working in resource-constrained environments.

Our framework and results are built on the foundation of shared and opensource materials, including tools like DeepLabCut [35], MMAction2 [41], and datasets such as OpenMonkeyChallenge [31] and PanAf500 [11]. This underscores the importance of making resources publicly available, especially in primatology, where data scarcity often impedes progress in AI-assisted methodologies. We strongly encourage researchers with large annotated video datasets to make them publicly accessible to foster interdisciplinary collaboration and further advancements in animal behavior research.

### 5.1 Challenges and Future Directions

While our results are promising, there are areas for improvement in both pose estimation and action recognition tasks.

#### Pose Estimation

Out-of-domain PCK metrics for pose estimation hovered just above 0.5, indicating that nearly half of the predicted keypoints were outside the acceptable range of the ground-truth coordinates. Accurate pose estimation is critical for downstream behavior classification. Future work could address this by finetuning the pose estimation model on the *behavior* dataset before pose extraction. Additionally, training on more specific datasets, such as OpenApePose [32], could improve performance. Techniques to reduce the domain gap between *pose* and *behavior* datasets [53] or leveraging pseudo-labels for semi-supervised learning [29, 54, 55] could also enhance generalization.

Interestingly, EfficientNet architectures performed worse than ResNet-152 in both within-domain and out-of-domain evaluations, contrary to prior results in animal pose estimation [34]. This discrepancy may stem from suboptimal hyperparameter tuning (e.g., fixed learning rate schedules instead of cosine decay) for EfficientNet models. Future studies should optimize hyperparameters individually for each architecture to fully explore their potential.

#### Behavior Recognition

While our skeleton-based pipeline achieved comparable results to previous studies, the overall accuracy remains relatively low, which could limit its practical deployment in the field. Comparisons to human-centric studies, where abundant datasets for both pose estimation and action recognition have led to higher performance [17], highlight the need for additional public datasets [28, 29, 56] to drive progress in AI-assisted animal behavior research.

From an algorithmic perspective, using keypoint detection as pose scoremaps rather than compressing them into (*x, y, c*) triplets could improve performance, particularly when pose predictions are less accurate [17]. Incorporating RGB-Pose dual-modality could further enhance classification accuracy, especially for behaviors with similar skeletal motion, such as ‘walking,’ ‘running,’ and ‘sitting on back.’

## 6 Conclusion

This study demonstrates the practical utility and relevance of skeleton-based action recognition approaches in animal behavior research. We hope the tools, methodologies, and insights presented here will inspire further applications of skeleton-based techniques to study a broader range of behaviors and animal species. Future advancements in pose estimation, action recognition, and dataset availability will undoubtedly enhance the impact of such approaches in ethology and beyond.

## 7 Acknowledgments

We extend our sincere gratitude to the team behind the Pan African Programme: ‘The Cultured Chimpanzee’, along with their partners, for granting us permission to use their data for this study. For access to the videos from the dataset, please reach out directly with the copyright holder Pan African Programme at http://panafrican.eva.mpg.de. In particular, we would like to thank H. Kuehl, C. Boesch, M. Arandjelovic, and P. Dieguez. Further acknowledgments go to: K. Corogenes, E. Normand, V. Vergnes, A. Meier, J. Lapuente, D. Dowd, S. Jones, V. Leinert, E. Wessling, H. Eshuis, K. Langergraber, S. Angedakin, S. Marrocoli, K. Dierks, T. C. Hicks, J-Hart, K. Lee, M. Murai and the team at Chimp&See.

The work that allowed for the collection of the PanAf500 dataset was made possible due to the generous support from the Max Planck Society, Max Planck Society Innovation Fund, and Heinz L. Krekeler. By extension, we also wish to thank: Foundation Ministre de la Recherche Scientifique, and Ministre des Eaux et Forêts in Cote d’Ivoire; Institut Congolais pour la Conservation de la Nature and Ministre de la Recherche Scientifique in DR Congo; Forestry Development Authority in Liberia; Direction des Eaux, Forêts Chasses et de la Conservation des Sols in Senegal; and Uganda National Council for Science and Technology, Uganda Wildlife Authority, and National Forestry Authority in Uganda.

In addition, we would like to thank the team at NCCR Evolving Language and in particular Guanghao You, for allowing us to use their computational platform.

## Appendix A

Prediction Comparison of Pose Estimation Models

**Fig. A1:**
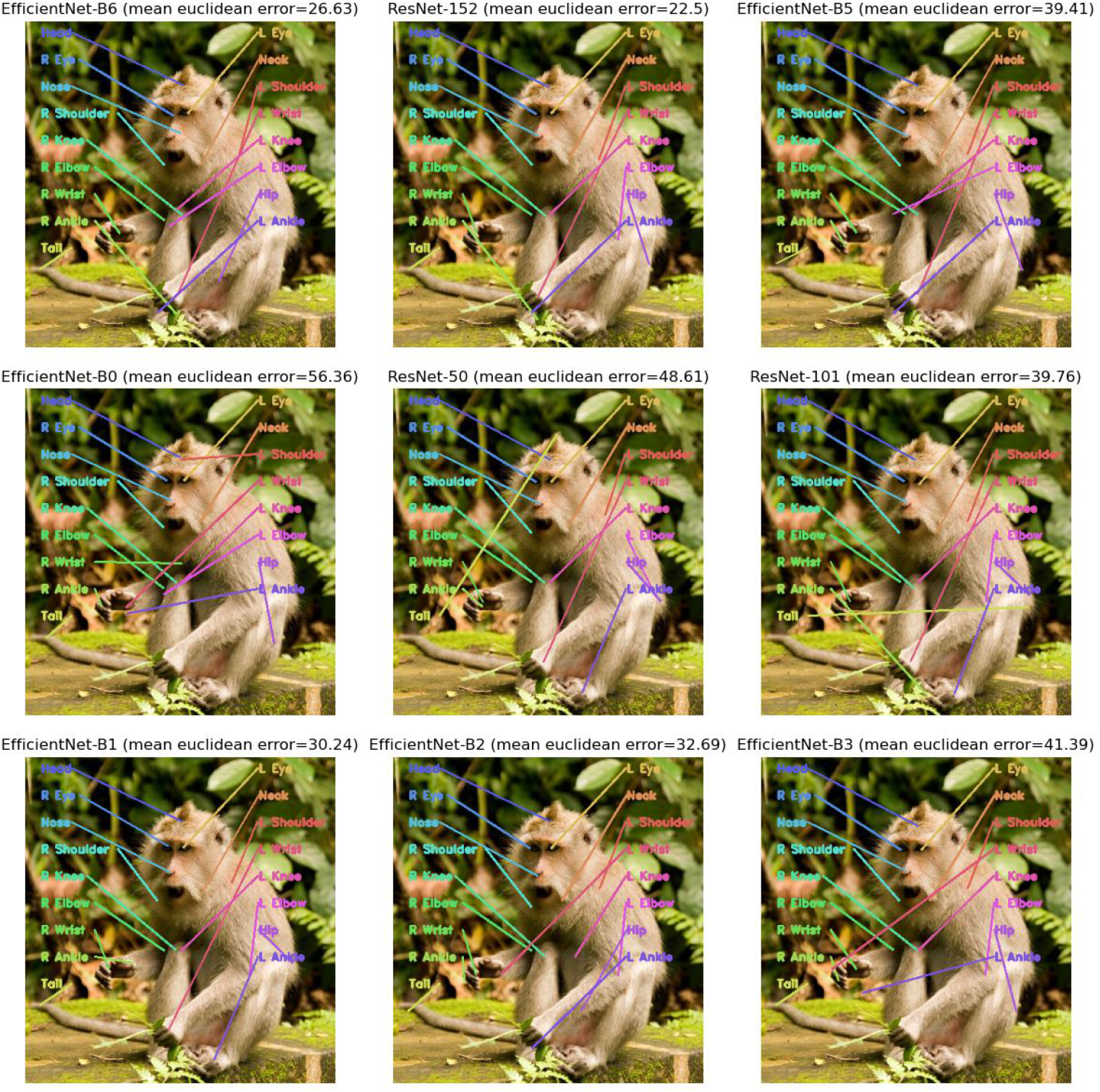
Prediction comparison of the nine models at test time. After 40,000 training iterations, the models’ test predictions are visually compared to one example of the test set. Note for example that i) ResNet-50 (*center*) wrongly predicts the top of the head as the tail’s position, ii) only three models can predict the left ankle’s position accurately (ResNet-50 (*center*), ResNet-101 (*center right*), and EfficientNet-B1 (*bottom left*)) and iii) no model correctly detects the left knee’s location.

## Appendix B

PCK Nasal Dorsum

**Fig. B2:**
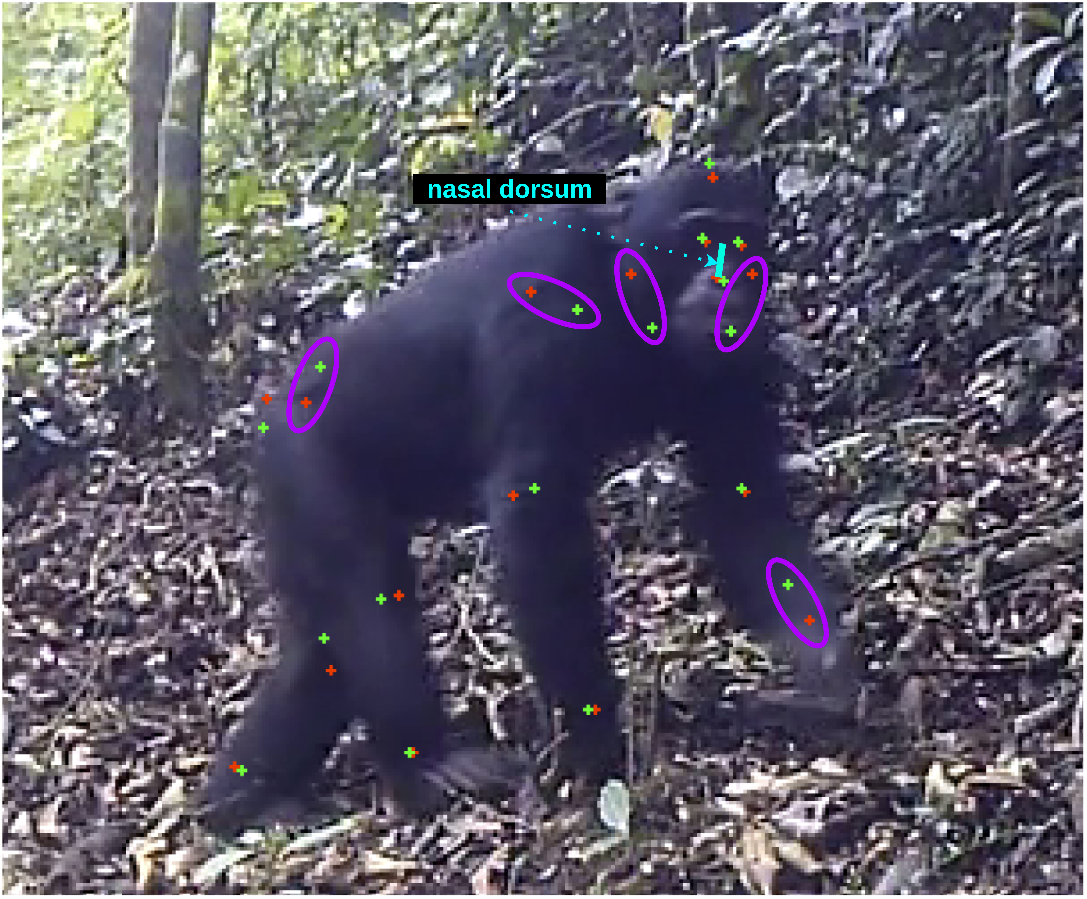
PCK nasal dorsum. The turquoise segment represents the length between the center of the eyes and the tip of the nose, i.e., the nasal dorsum. Any model prediction (represented in green) that falls within this distance of the ground-truth location (indicated in red) is considered as detected. In this case, all keypoints are detected except for the shoulders, neck, left wrist, and the hip (circled in purple). Hence, for this image, the detection rate would be 12*/*17 = 0.706 = 70.56%.

## Appendix C

NMER by Families, Species and Keypoints

**Fig. C3:**
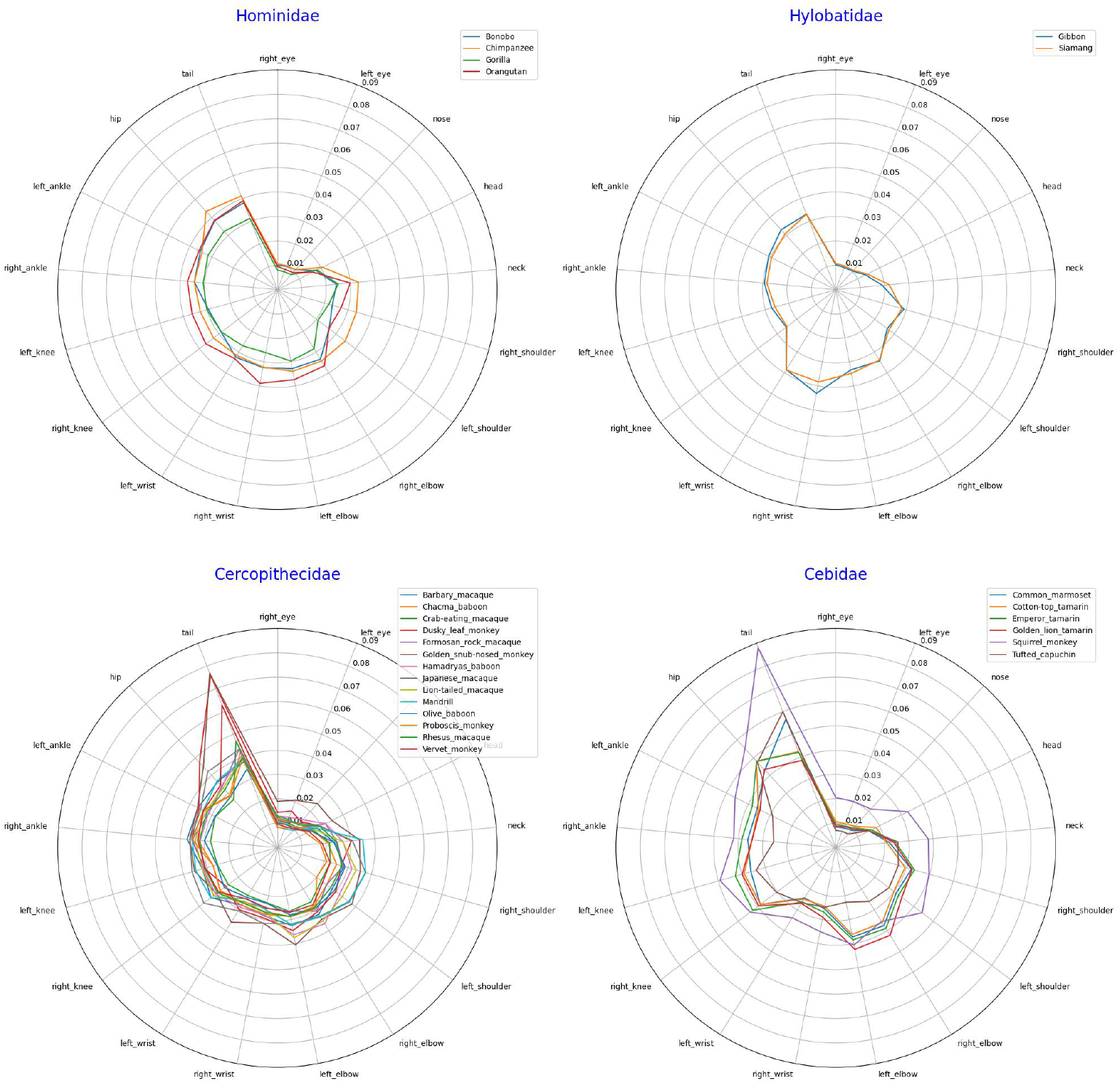
Normalized error rate by families, species and keypoints. For all OMC images at test time, we visualize the normalized error rate (NMER) for each species.

## Appendix D

Examples of Elements of the ASBAR GUI

**Fig. D4:**
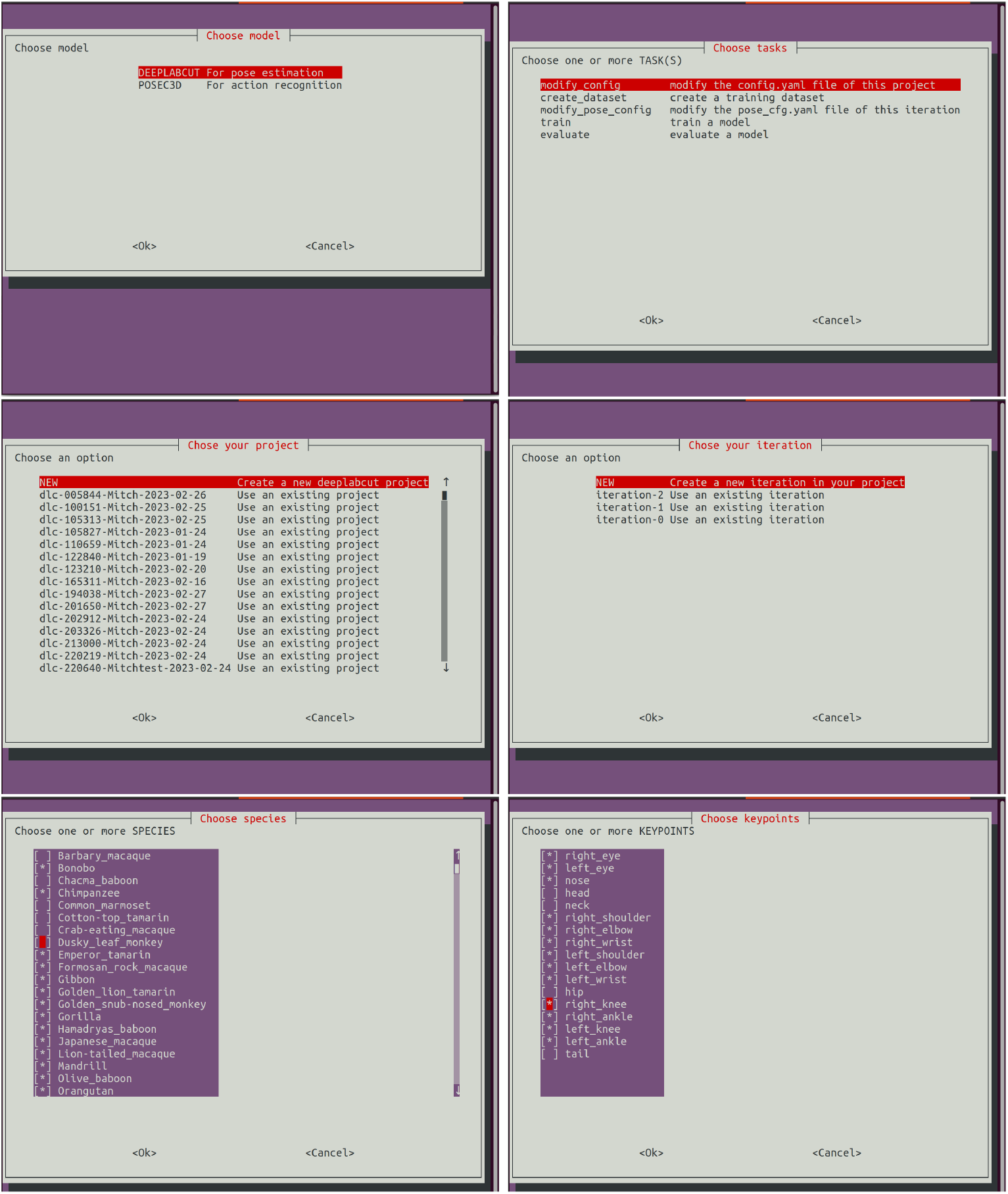
Examples of UI elements of the ASBAR graphical user interface. The GUI is terminal-based and therefore can be rendered even when accessed on a distant machine, such as a cloud-based platform or a remote high-performance computer.

